# mTORC1-Plin3 pathway is essential to activate lipophagy and protects against hepatosteatosis

**DOI:** 10.1101/812990

**Authors:** Marina Garcia-Macia, Adrián Santos-Ledo, Jack Leslie, Hanna Paish, Abigail Watson, Lee Borthwick, Jelena Mann, Fiona Oakley, Viktor I. Korolchuk, Derek A. Mann

**Affiliations:** Institute of Cellular Medicine, Newcastle University, Newcastle Upon Tyne, UK; Instituto de Investigación Biomédica de Salamanca (IBSAL), Universidad de Salamanca, Institute of Functional Biology and Genomics, CSIC, Zacarias Gonzalez, 2, 37007 Salamanca, Spain; Institute of Genetic Medicine, Newcastle University, Newcastle Upon Tyne, UK; Institute for Cellular and Molecular Biosciences, Newcastle University, Newcastle Upon Tyne, UK

**Keywords:** lipophagy, Plin3, mTORC1, autophagy, hepatocytes

## Abstract

During postprandial state, the liver is exposed to high levels of dietary fatty acids (FAs) and carbohydrates. FAs are re-esterified into triglycerides, which can be stored in lipid droplets (LDs) in the liver. Aberrant accumulation of LDs can lead to diseases such as alcoholic liver disease and non-alcoholic fatty liver disease, the latter being the most common liver pathology in western countries. Improved understanding of LD biology has potential to unlock new treatments for these liver diseases. The Perilipin (Plin) family is the group of proteins that coat LDs, controlling their biogenesis, stabilization, and preventing their degradation. Recent studies have revealed that autophagy is involved in LD degradation and, therefore, may be crucial to avoid lipid accumulation. Here, we show that a phosphorylated form of Plin3 is required for selective degradation of LDs in fibroblasts, primary hepatocytes and human liver slices. We demonstrate that oleic acid treatment induces the recruitment of the autophagy machinery to the surface of LDs. When Plin3 is silenced, this recruitment is suppressed resulting in accumulation of lipid. Plin3 pulldowns revealed interactions with the autophagy initiator proteins Fip200 and Atg16L indicating that Plin3 may function as a docking protein involved in lipophagy activation. Of particular importance, we define Plin3 as a substrate for mTORC1-dependent phosphorylation and show that this event is decisive for lipophagy. Our study therefore reveals that Plin3, and its phosphorylation by mTORC1, is crucial for degradation of LDs by autophagy. We propose that stimulating this pathway to enhance LD autophagy in hepatocytes will help protect the liver from lipid-mediated toxicity thus offering new therapeutic opportunities in human steatotic liver diseases.

## INTRODUCTION

After food consumption, also known as a postpandrial state, the liver handles high levels of nutrients using some as a fuel (mostly lipids), whilst also mediating their mobilization to the whole organism or storage in the form of glycogen (Matikainen, Manttari, Westerbacka, Vehkavaara, Lundbom, Yki-Jarvinen and Taskinen, 2007). Caloric excess and sedentary lifestyle have led to a global epidemic of obesity and metabolic syndrome, which in approximately one third of the population causes non-alcoholic fatty liver disease (NAFLD) (Anstee, Reeves, Kotsiliti, Govaere and Heikenwalder, 2019). NAFLD is the most frequent liver pathology in western countries (Younossi, Anstee, Marietti, Hardy, Henry, Eslam, George and Bugianesi, 2018) and in some individuals can progress to advanced fibrosis, cirrhosis and cancer (Mortality and Causes of Death, 2016, Younossi, Anstee, Marietti, Hardy, Henry, Eslam, George and Bugianesi, 2018). Unfortunately, there are no efficient treatments to prevent or ameliorate NAFLD. It is reported that diet and exercise have a positive impact on NAFLD (Romero-Gomez, Zelber-Sagi and Trenell, 2017). However, the exact links between lifestyle and hepatocyte biology are still poorly understood.

Under stress situations such as obesity or alcohol-induced damage, hepatocytes accumulate fat in lipid droplets (LDs), which contributes to the development of hepatosteatosis (Gluchowski, Becuwe, Walther and Farese, 2017). LDs are dynamic metabolic stations (Greenberg, Kraemer, Soni, Jedrychowski, Yan, Graham, Bowman and Mansur, 2011), formed by a core of neutral lipids surrounded by a phospholipid membrane and proteins, of which members of the perilipin (Plin) family are amongst the most important. Perilipins are a family of proteins that coat LDs, controlling their biogenesis, stabilization, and preventing their degradation. Plin2 and Plin3 are expressed in hepatocytes (Pawella, Hashani, Eiteneuer, Renner, Bartenschlager, Schirmacher and Straub, 2014), which makes them a potential target to promote selective degradation of lipids and limit LD accumulation (Kaushik and Cuervo, 2015).

LD catabolism can be initiated downstream of two distinct stimuli: nutrient deprivation and acute LD overload. Both scenarios have in common an increase in autophagy activity (Niso-Santano et al., 2015, Singh, Kaushik, Wang, Xiang, Novak, Komatsu, Tanaka, Cuervo and Czaja, 2009). Autophagy is the cellular clearing and recycling program that degrades unwanted cytoplasmic content within lysosomes (Singh, Kaushik, Wang, Xiang, Novak, Komatsu, Tanaka, Cuervo and Czaja, 2009, Mizushima, 2009, Singh et al., 2009). This process takes place in all cells and is further induced during cellular stress. Besides its crucial role as a quality control mechanism, autophagy is also involved in a host of other functions such as growth and differentiation (Mizushima, 2009), metabolic regulation and as an alternative energy source (Singh, Kaushik, Wang, Xiang, Novak, Komatsu, Tanaka, Cuervo and Czaja, 2009, Singh et al., 2009).

To date, at least two mechanisms for LD degradation through autophagy have been described. One involves chaperone-mediated autophagy (CMA) where the LD perilipin Plin2, also known as ADRP or Adipophilin, and probably also Plin3, known as Tip47, binds to the lysosome protein Lamp2a and mediates LD uncoating before autophagosome engulfment (Kaushik and Cuervo, 2015, Kaushik and Cuervo, 2016). The other mechanism involves macroautophagy (hereafter autophagy) where LDs are engulfed in the cytoplasm and then delivered to the lysosome via autophagosome–lysosome fusion. This latter type of autophagy is called lipophagy (Singh and Cuervo, 2012). Selective autophagic degradation of cellular organelles, including mitochondria (Ashrafi and Schwarz, 2013), peroxisomes (Hutchins, Veenhuis and Klionsky, 1999), lysosomes (Hung, Chen, Yang and Yang, 2013), endoplasmic reticulum (ER) (Smith and Wilkinson, 2018) and the nucleus (Nakatogawa and Mochida, 2015) is mediated by specific receptor proteins. For example, BNIP3, NIX, OPTN, NDP62 and TAX1BP1 have been implicated in autophagic degradation of mitochondria (mitophagy) (Khaminets, Behl and Dikic, 2016). However, to our knowledge, specific receptors for lipophagy remain unknown. From the LD-resident proteins, small GTPases of Rab family may potentially assist with the recruitment of the autophagic machinery to the LD (Schroeder, Schulze, Weller, Sletten, Casey and McNiven, 2015), but perilipins are the only proteins specific to LDs (Kaushik and Cuervo, 2015). Both Plin2 and Plin3 need to be degraded, following phosphorylation, for an efficient lipophagy (Kaushik and Cuervo, 2016). Interestingly, Plin3 is found in CMA-deficient lysosomes in fed state which indicates a different degradation pathway and function for Plin3 compared to Plin2 which is only found in CMA-proficient lysosomes (Kaushik and Cuervo, 2015). Intriguingly, this observation also suggests that, unlike bulk autophagy, lipophagy may not be suppressed in response to nutrient abundance.

The key cellular sensor of nutrients is the mammalian target of Rapamycin (mTOR)(Rabanal-Ruiz and Korolchuk, 2018). mTOR is the catalytic subunit of two distinct complexes, called mTOR Complex 1 (mTORC1) and 2 (mTORC2). These two complexes have unique protein components and respond to different cellular signalling events with distinct outcomes (Laplante and Sabatini, 2009). In nutrient replete condition mTORC1 promotes cellular anabolism whilst suppressing autophagy. mTORC1 has also been shown to promote the formation of LDs (Mensah, Goberdhan and Wilson, 2017). Interestingly, simultaneously with mTOR activation, the high levels of nutrients including lipids activate lipophagy to prevent hepatic lipotoxicity in the liver (Singh, Kaushik, Wang, Xiang, Novak, Komatsu, Tanaka, Cuervo and Czaja, 2009). The exact mechanisms behind these seemingly contradictory observations remain unresolved.

To address this issue, we have emulated acute lipid overload with oleic acid treatment in different experimental models (*in vitro* and *ex vivo*) and analysed mechanisms allowing for simultaneous activation of mTOR and lipophagy. We identified phosphorylated Plin3 as a specific receptor for lipophagy after acute lipid overload, and mTOR as the kinase responsible for this phosphorylation both *in vitro* and in human tissue. Our findings provide an important insight into the mechanisms of energy utilization in the liver and identify potential targets for the selective degradation of LDs which may lead to future therapeutic interventions in different diseases including obesity, atherosclerosis and NAFLD.

## RESULTS

### Oleic acid treatment induces lipophagy and selective degradation of Plin3

Cells treated with fatty acids (FAs) provide an excellent tool to emulate the lipid overload characteristic of the postprandial state and, at high doses, they are used to simulate disease conditions such as those found in NAFLD (Alkhatatbeh, Lincz and Thorne, 2016, Liu, Liao, Chen and Han, 2019). Oleic acid (OA) treatment was chosen as a model to induce LD formation (Rohwedder, Zhang, Rudge and Wakelam, 2014, Lagrutta, Montero-Villegas, Layerenza, Sisti, Garcia de Bravo and Ves-Losada, 2017). This monosaturated FA is able to support adaptative mechanisms, including triglyceride synthesis, to prevent cellular stress (Fu, Watkins and Hotamisligil, 2012). It is known that autophagy is also triggered following OA treatment (Fig. S1A) (Niso-Santano et al., 2015), which is not a response to cellular stress. Thus, the first aim was to identify whether OA triggered autophagy can selectively degrade LDs. For that purpose, NIH-3T3 fibroblasts and/or primary hepatocytes cultured with OA were treated with lysosomal inhibitors prior to determination of autophagy flux and analysis of levels of autophagy cargo proteins (Klionsky et al., 2016). In addition, two complementary approaches were used to analyse lipophagy activity: protein analysis from isolated LDs by western blot and protein colocalization by immunofluorescence (Fig. 1A).

**Figure 1:**
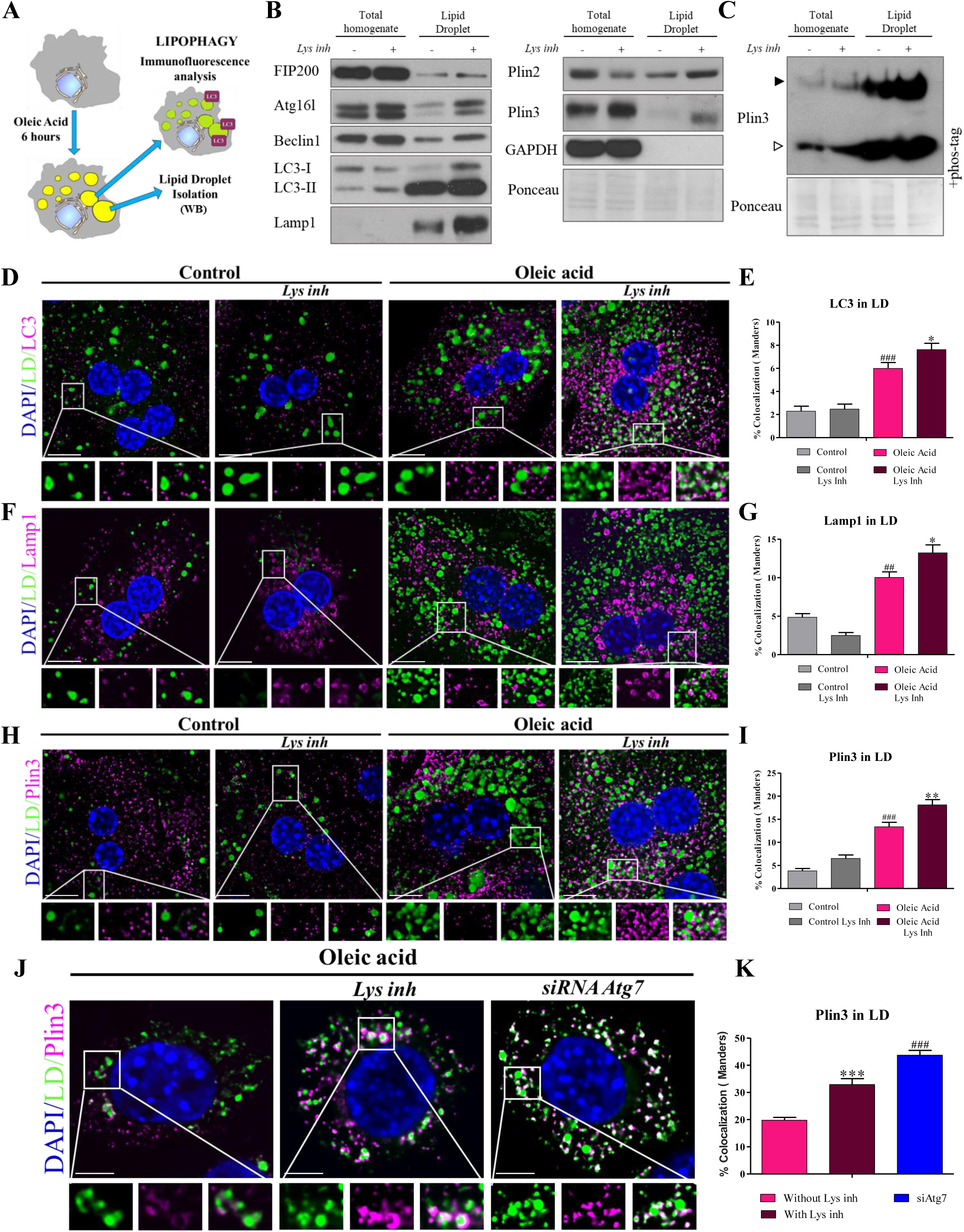
Oleic acid treatment induces lipophagy and selective degradation of Plin3. A) Depiction of the methodological approach: Oleic acid (OA) treatment induces lipid droplet (LD) formation. Lipophagy is analysed by immunofluorescence and western blotting in homogenates and isolated LDs. B) Autophagy pathway proteins and perilipins expression in total homogenates and LD isolations at 6 hours of OA treatment with or without lysosomal inhibitors (Lys inh) to test autophagy flux. LC3 flux in total homogenates shows active autophagy, supporting selective degradation of LDs. All autophagy proteins tested accumulate in the LD isolations after lysosomal inhibitors treatment. Plin3 is selectively degraded by autophagy, shown both in homogenates and isolated LDs. C) Total homogenates and LDs were subjected to Phos-tag gel electrophoresis and immunoblotted for Plin3. Phosphorylated Plin3 (p-Plin3: black arrowhead) is mainly found in LDs. Phosphorylation is increased after blocking autophagy with lysosomal inhibitors. D-G) LC3 and Lamp1 (magenta) recruitment to LDs (green) is enhanced after blocking autophagy in the OA-treated primary hepatocytes (D and F, quantified in E and G). H, I) Plin3 (magenta) recruitment to LDs (green) is also enhanced after blocking autophagy (H, quantified in I). J, K) Plin3 recruitment is also enhanced after blocking autophagy using Atg7 knockdown in OA-treated NIH-3T3 cells. Scale bar: 20μm. Bars are mean ± SEM. *p < 0.05, **p < 0.01, and ***p < 0.001 (differences caused by lysosomal inhibitors treatment), ^#^p<0.05, ^##^p<0.01, and ^###^p<0.001 (differences caused by treatment).

Proteins involved in autophagy initiation (Fip200 (also known as Rb1cc1), Atg16l and Beclin 1 were mildly increased after blocking lysosomal fusion in total cell lysates and in LD fractions (Fig. 1B). At the same time, LC3-II and Lamp1, autophagosomal and lysosomal markers respectively, were strongly accumulated in LDs indicating a substantive induction of lipophagy activity after OA treatment (Fig. 1B). Between the two most abundant Perilipins in hepatocytes, Plin3 was more abundant after lysosomal inhibition, this suggesting that Plin3 is a preferential target for lysosomal degradation compared to Plin2 (Fig. 1B). Plin3 degradation depends on its phosphorylation, this enabling the autophagic machinery to access LDs (Kaushik and Cuervo, 2015). Therefore, electrophoresis with Phos-tag was used to facilitate resolution of phosphorylated variants of Plin3 (Kaushik and Cuervo, 2016). This analysis revealed an increase in the phosphorylated form of Plin3 (upper band) in LDs after treatment with lysosomal inhibitors, hence phosphorylated Plin3 is likely to be a direct target of autophagy degradation (Fig. 1C).

To confirm the previous results, we co-stained hepatocytes with LC3 (Fig. 1D, E) and Lamp1 (Fig. 1F, G) together with BODIPY, a dye that stains neutral lipids in LDs and performed these stains in the presence and absence of OA treatment. Co-localization of LC3 and Lamp1 with LDs in hepatocytes increased after OA treatment (Fig. 1D-G, p<0.01). Accumulation around LDs was further increased after the treatment with lysosomal inhibitors (Fig. 1D-G, p<0.001). A similar pattern was observed for Plin3 where OA treatment strongly increased its autophagic flux (Fig. 1H and I, p<0.01). To verify the degradation of Plin3 by autophagy, we silenced *Atg7* (an essential autophagy gene, Fig. S1B). Plin3 was further accumulated on the LD surface after *Atg7* silencing (Fig. 1J-K, p<0.001), supporting the idea of Plin3 as a selective cargo for OA-induced lipophagy. Based on these data we conclude that lipid overload, emulated by the OA treatment, induces lipophagy and selective degradation of Plin3.

### Plin3 promotes recruitment of autophagy machinery to LDs

The role of Plin3 in lipophagy was interrogated by siRNA-mediated knock-down. In response to loss of Plin3 expression we observed no effect on expression of the macroautophagy core machinery (Fig. 2A-D). However, the recruitment of LC3 (Fig. 2E and F) and Lamp1 (Fig. 2G and H) to LD was strongly suppressed after Plin3 silencing (Fig. 2E-H, p<0.01). To further determine if Plin3 functions as a key component of the autophagy machinery in LDs, we probed for Fip200 and Atg16l in the immunoprecipitated endogenous Plin3 fractions. The association of these autophagy initiation proteins with Plin3 was observed to be increased following OA treatment (Fig. 2I), this suggestive of Plin3 physically associating with the autophagy machinery in response to OA. Interestingly, this co-immunoprecipitation was abolished by rapamycin treatment (see Fig. 4). Fip200 is recruited to the sites of autophagosome biogenesis, preceding and facilitating the recruitment of other ATG proteins (Ktistakis and Tooze, 2016). Hence, we were interested to determine next if Plin3 influences key downstream partners of Fip200 that are required for activation of autophagy, in particular Beclin 1. To this end, endogenous Fip200 pulldowns were examined using cells cultured with or without OA treatment and Plin3 silencing (Fig. S1C). Interaction of Fip200 and Beclin 1 was increased after OA treatment, as expected as lipophagy was also increased, however the interaction was suppressed in Plin3 silenced cells (Fig. 2J). These results indicate a crucial role for Plin3 in the recruitment of autophagic machinery to LDs. As such, we propose Plin3 acts as a previously unrecognised lipophagy receptor that is required for assembling autophagic components in LD.

**Figure 2:**
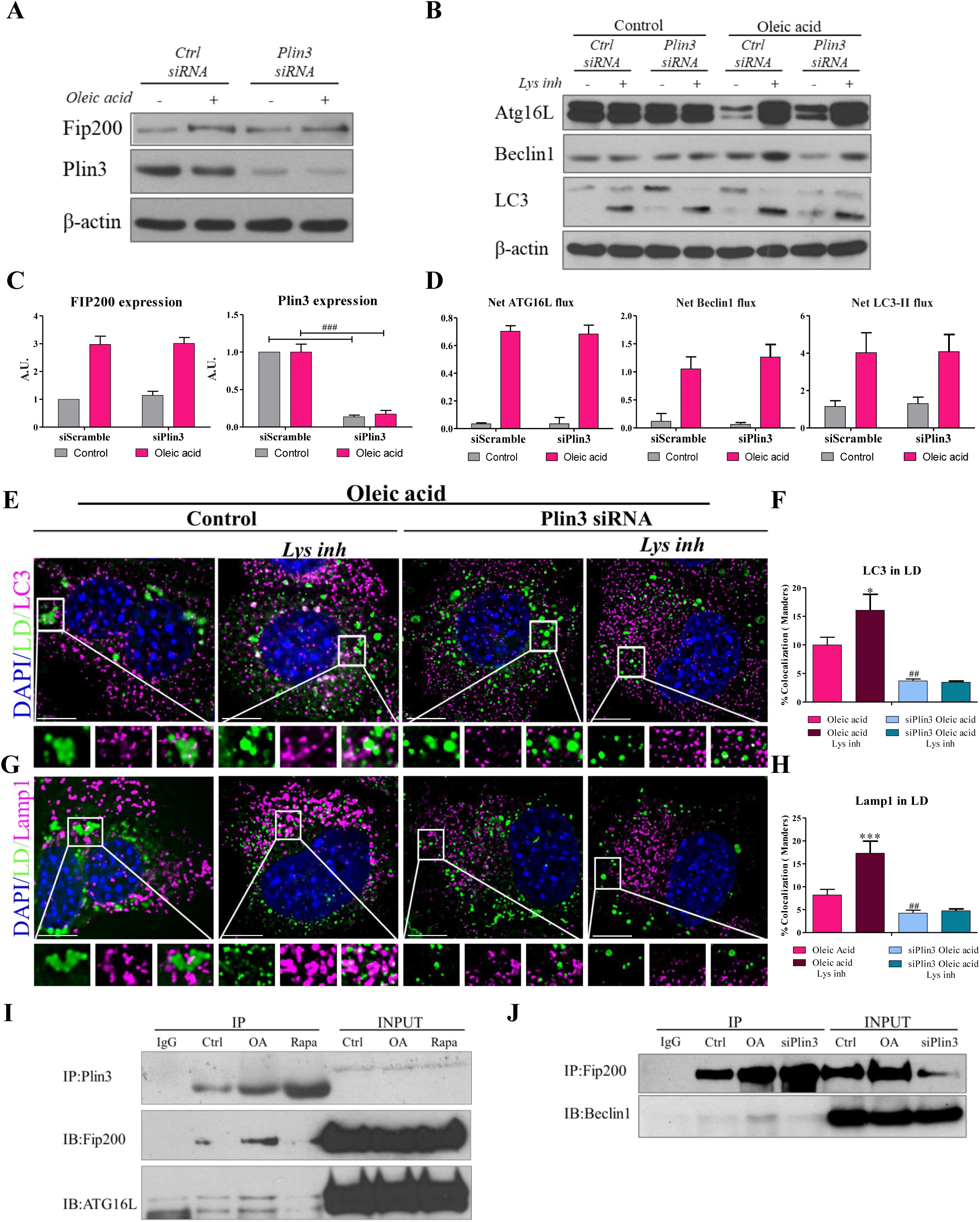
Plin3 docks autophagy machinery to LDs. A-D) Levels of Plin3 and autophagy pathway proteins in total homogenates from control and Plin3 silenced cells with and without OA and with and without treatment with lysosomal inhibitors. There are no differences in the expression of autophagy proteins between control and silenced cells (quantified in C (protein expression) and D (protein flux). E-H) LC3 and Lamp1 (magenta) recruitment to LDs (green) is reduced after silencing Plin3 in NIH-3T3 fibroblasts treated with OA (showed in C and E, quantified in D and F) indicating impairment of lipophagy in siPlin3 cells. I) Binding between Plin3 and the autophagy initiator proteins Fip200 and ATG16l assessed by the co-immunoprecipitation with endogenous Plin3 is increased after OA treatment. J) OA treatment increases the binding between Fip200 and Beclin, this binding is reduced after silencing Plin3. Scale bar: 20μm. Bars are mean ± SEM. *p < 0.05, **p < 0.01, and ***p < 0.001 (differences caused by lysosomal inhibitors treatment), ^#^p<0.05, ^##^p<0.01, and ^###^p<0.001 (differences caused by treatment).

### mTORC1 associates with LDs

Precise regulation of LD biology relies on multiples cues controlled by nutrient levels (Nguyen and Olzmann, 2019). mTORC1 is activated during overfeeding (Zoncu, Efeyan and Sabatini, 2011) and also after acute lipid overload (Fig. S2A). It is also reported that mTORC1 mediates lipid accumulation in fat cells (Zoncu, Efeyan and Sabatini, 2011). Levels of total and phosphorylated mTOR (Ser2448) were increased in LDs upon blockade of lysosomal activity (Fig. 3A). Raptor, but not Rictor, components of mTORC1 and mTORC2 respectively, was slightly increased in LDs after treatment with lysosomal inhibitors (Fig. 3A). Similarly, Rheb, the master activator of mTORC1, was found to accumulate in LD fractions after lysosomal inhibition (Fig. 3A). mTORC1 recruitment to the surface of the lysosome, which is required for its activation, is mediated by a heterodimeric complex of Rag GTPases, Rag A/B or Rag C/D (Sancak, Peterson, Shaul, Lindquist, Thoreen, Bar-Peled and Sabatini, 2008, Sancak, Bar-Peled, Zoncu, Markhard, Nada and Sabatini, 2010). Rag GTPases were also found in LD accumulated in response to lysosomal inhibitors (Fig. 3A). TSC1, a component of the TSC complex which acts as a mTOR inhibitor, was absent in LDs. These data support the presence of the machinery required for mTORC1 activation in LDs and describe an unexpected cellular localization of mTORC1.

**Figure 3:**
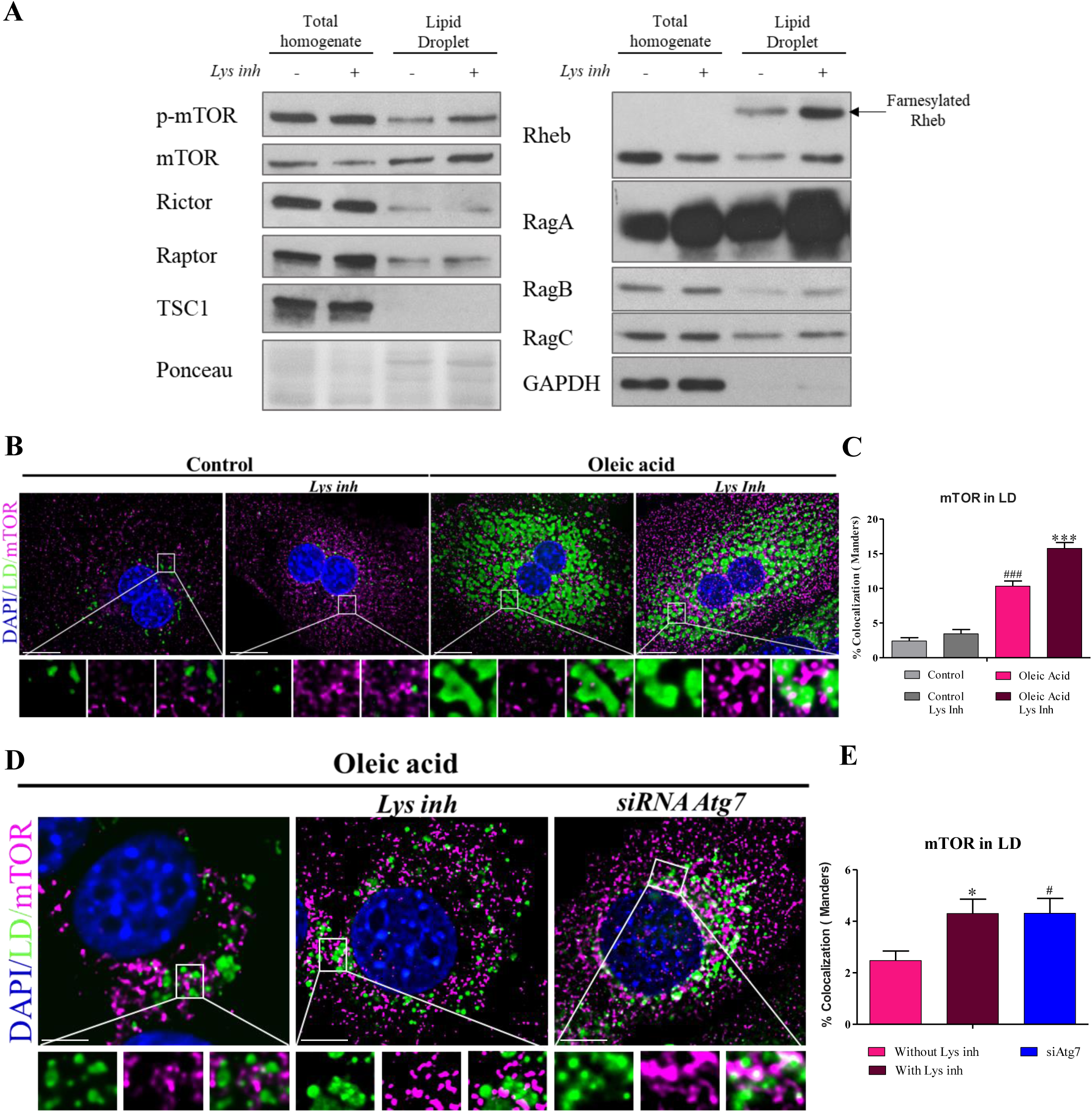
mTORC1 machinery is present in LDs. A) Levels of mTOR pathway proteins in total homogenates and LD isolations at 6 hours of OA treatment. Active mTOR, Raptor, Rheb and RagA, B and C, but not Rictor or TSC1, accumulate in LDs after lysosomal inhibition. B-C) mTOR (magenta) recruitment to LDs (green) is enhanced after blocking autophagy in the OA-treated primary hepatocytes (B, quantified in C). D-E) mTOR recruitment is also enhanced after blocking autophagy through Atg7 silencing in OA-treated NIH-3T3 cells. (D quantified in E). Scale bar: 20μm. Bars are mean ± SEM. *p < 0.05, **p < 0.01, and ***p < 0.001 (differences caused by lysosomal inhibitors treatment), ^#^p<0.05, ^##^p<0.01, and ^###^p<0.001 (differences caused by treatment)

To further confirm that mTOR is present in LDs, co-localization with BODIPY was performed. As predicted, recruitment of mTOR to LDs was enhanced after OA treatment (Fig. 3B and C, p<0.01), and levels of mTOR further increased after blocking lysosomal function (Fig. 3B and C, p<0.001). This indicated active lysosomal degradation of mTOR from the LD surface. To determine whether degradation of mTOR was mediated by autophagy, Atg7 was silenced in NIH-3T3 cells. As expected, mTOR accumulated in LDs after blocking autophagy by Atg7 silencing and to the same extent as observed upon treatment with lysosomal inhibitors (Fig. 3D and F, p<0.05).

Together these data indicate that mTORC1 is present on LDs, potentially anchored to their surface by Rag GTPases. Moreover, mTOR is present on LDs in its active, phosphorylated form which is consistent with the presence of its master activator Rheb. Interestingly, components of mTORC1 in LDs appear to undergo lysosomal degradation mediated by autophagy.

### Role of Plin3 in lipophagy is controlled by mTORC1

To explore the function of mTORC1, rapamycin treatment was used to block its activity (Fig. 4A, Fig. S2B). After rapamycin treatment, mTOR and p-mTOR were barely detectable in LDs, while mTOR levels were unaffected in the total homogenates (Fig. 4B). Similarly, farnesylated Rheb was also decreased in LDs after rapamycin treatment and lost the autophagy flux effect (Fig. 4B). As mTORC1 is the main autophagy inhibitor, levels of autophagy proteins were tested in LDs. Autophagy initiator proteins (Fip200, Atg16l and Beclin 1) were increased in the total homogenates from cells treated with rapamycin, as expected. Strikingly, their levels decreased in LDs from cells treated with rapamycin (Fig. 4B). Similarly, the recruitment of LC3-II and Lamp1 to LDs was also inhibited after rapamycin treatment (Fig. 4B). These results suggest that mTOR inhibition by rapamycin disturbs recruitment of the autophagy machinery to LDs. This conclusion was corroborated by colocalization analyses of LC3 and Lamp1 with BODIPY (LDs) in primary hepatocytes. In OA treated cells, LC3 localization to LDs was decreased after exposure to rapamycin (Fig. 4C and D, p<0.05) whilst Lamp1 colocalization with BODIPY was almost absent in response to rapamycin treatment (Fig. 4E and F, p<0.001). To test if mTORC1 activity affects Plin3, Plin3 colocalization with BODIPY was also tested. Rapamycin treatment enhanced Plin3 accumulation on the surface of LDs (Fig. 4G and H, p<0.05). Similar results were obtained when mTORC1 was silenced in NIH-3T3 cells (Fig. S2C-H).

**Figure 4:**
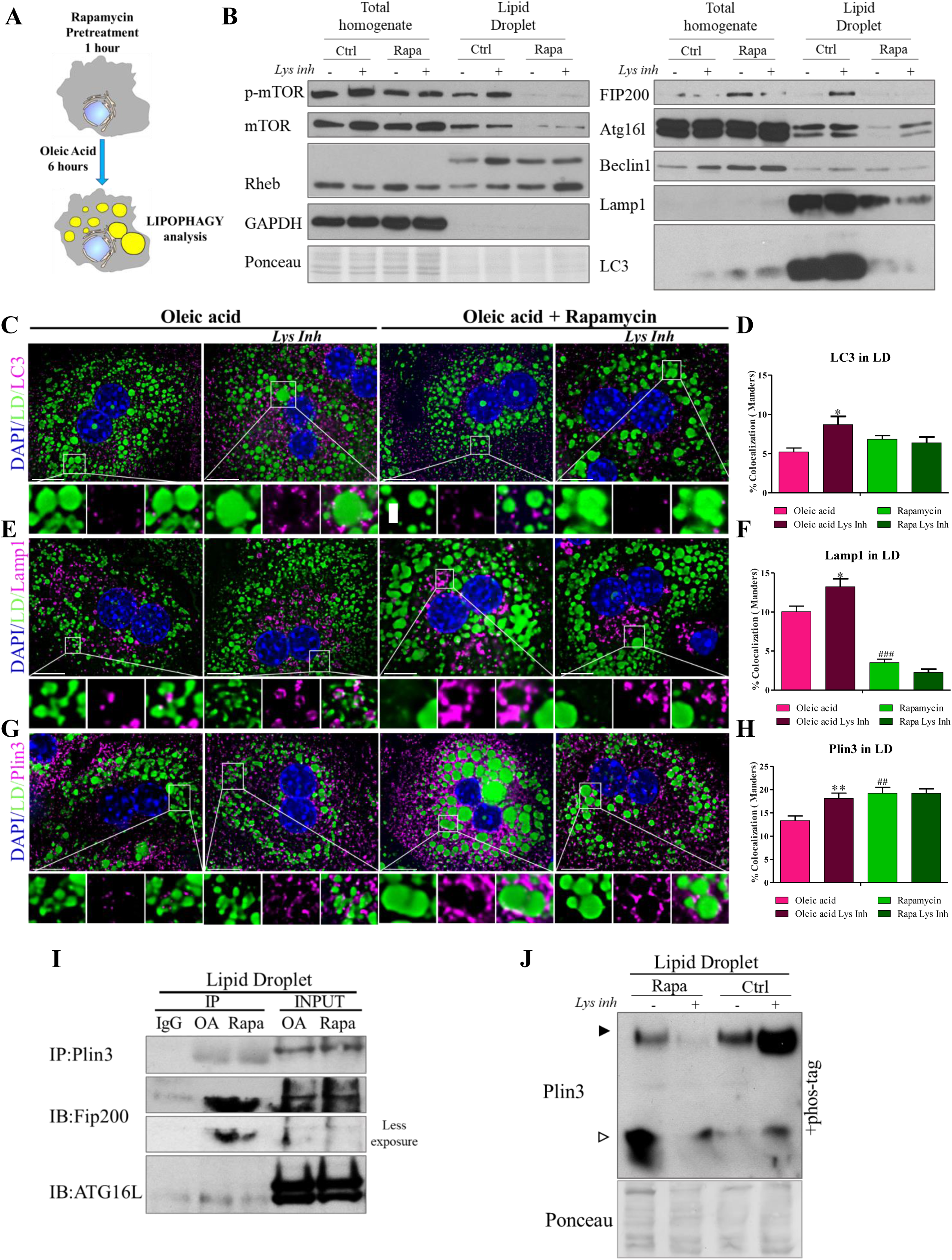
The role of Plin3 on LDs is controlled by mTORC1 activity. A) Depiction of the methodological approach: 1h rapamycin treatment before the induction of LD formation by OA treatment. Lipophagy is tested as previously described. B) Autophagy and mTORC1 pathway protein levels in total homogenates and LD isolations at 6 hours of OA treatment with and without rapamycin pretreatment. Note that levels of autophagy proteins are decreased after rapamycin treatment; active mTOR and Rheb are also reduced when mTORC1 activity is inhibited. C-H) LC3 and Lamp1 (magenta) recruitment to LDs (green) is impaired after blocking mTORC1 activity (C and E, quantified in D and F). However, Plin3 (magenta) recruitment to LDs (green) is enhanced after rapamycin pretreatment in OA-treated primary hepatocytes (G, quantified in H). I) Immunoprecipitation of endogenous Plin3 from LD isolations indicates its reduced binding to Fip200 and ATG16l after rapamycin treatment. J) Plin3 phosphorylation in LDs is diminished after rapamycin treatment. Scale bar: 20μm. Bars are mean ± SEM. *p < 0.05, **p < 0.01, and ***p < 0.001 (differences caused by lysosomal inhibitors treatment), ^#^p<0.05, ^##^p<0.01, and ^###^p<0.001 (differences caused by treatment)

As mTORC1 controls autophagic turnover of Plin3, we next determined if it also influences Plin3-mediated recruitment of autophagy machinery to LDs. We therefore immunoprecipitated endogenous Plin3 from LDs isolated from OA-treated cells that had been pre-treated with rapamycin. Both Fip200 and Atg16l were co-precipitated with Plin3 on LDs formed in response to OA, but of note this interaction was diminished in rapamycin pretreated cells (Fig. 4I). Hence, mTORC1 inhibition is detrimental for the binding of Plin3 to autophagy activators. Finally, we used Phos-tag approach to test the effect of rapamycin on Plin3 phosphorylation. mTORC1 inhibition reduced the levels of phosphorylated Plin3 and the flux effect was also abolished (Fig. 4J). Collectively our results indicate that Plin3 phosphorylation is a key event in docking autophagic machinery to LDs during lipophagy and that mTOR is the kinase behind this crucial phosphorylation.

### Inhibition of the mTORC1-Plin3 axis suppresses fuelling of the mitochondria by fatty acids

Lipophagy is the most effective way to mobilize fatty acids (FAs), thus preventing FA-mediated toxicity (Nguyen, Louie, Daniele, Tran, Dillin, Zoncu, Nomura and Olzmann, 2017). In order to understand the impact of the modulation of autophagy on cell metabolism, the oxygen consumption (OCR) was measured by Seahorse (Fig. 5A). Oleic acid-treated primary hepatocytes displayed higher mitochondrial and maximal respiration than hepatocytes under basal conditions (Fig. 5B and C). This increase in respiration was reversed when CPT1, an enzyme involved in the transport of FAs to the mitochondria, was inhibited using etomoxir (Fig. 5B and C), when autophagy was blocked (using lysosomal inhibitors), or when mTOR was inactivated (with rapamycin) (Fig. 5B and C, p<0.001). Concomitantly, ATP production was also increased after OA treatment, this effect being supressed when etomoxir, lysosomal inhibitors or rapamycin were applied (Fig. 5D, p<0.05). Similar results were obtained in NIH-3T3 cells where OA treatment increased oxygen consumption and ATP production (Fig. 5E, F, G, Fig. S3). Interestingly, Plin3 knockdown prevented an energetic response to OA and unlike the siScramble control, cells that were treated with Plin3 siRNA did not increase their maximal respiration or ATP production (Fig. 5E, F and G, Fig. S3). We conclude that Plin3 is crucial for the mobilization of fat for energy generation. Hence, the mTOR-Plin3 mechanism identified in this study is not only crucial to initiate lipophagy but is also paramount for successful energy production and avoidance of lipotoxicity after lipid overload.

**Figure 5.**
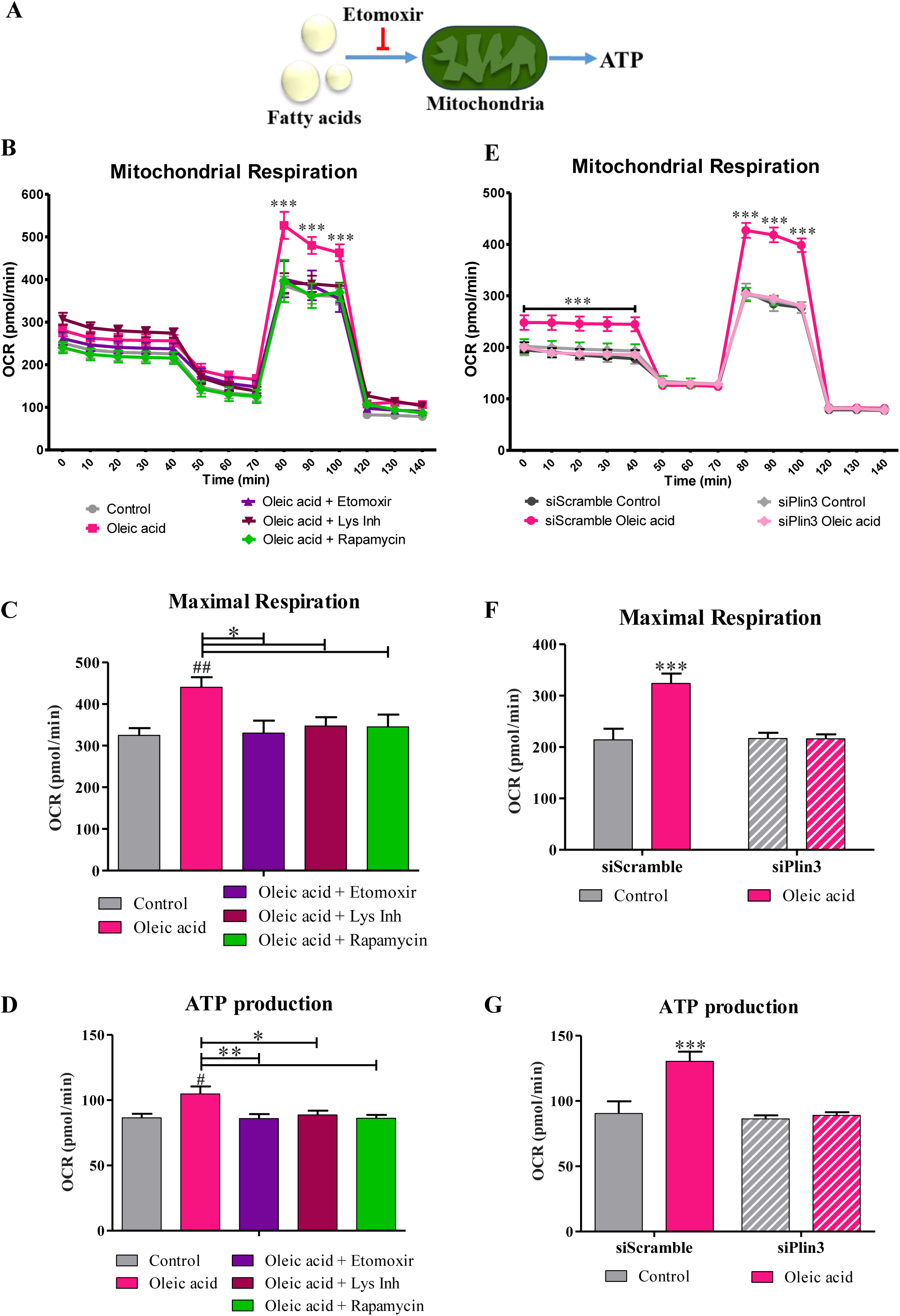
Lipophagy produces fuel for mitochondria which is inhibited upon mTORC1 inactivation. A) Schematic diagram of the metabolic function of lipophagy. Fatty acids (FAs) produced by lysosomes are used by mitochondria to produce ATP. Etomoxir inhibits the CPT1a, which transports FAs to the mitochondria. B) Mitochondrial respiration in primary hepatocytes under basal condition and after 6 hours of OA treatment with and without etomoxir (100nM), rapamycin or lysosomal inhibitors pretreatment. OA treatment increases mitochondrial respiration in hepatocytes in basal state, which is prevented in the presence of mitochondrial FA transport (etomoxir), autophagy (Lys inh) or mTORC1 (rapamycin) inhibitors. C and D) Maximal respiration and ATP production are increased in hepatocytes in response to OA treatment, which is suppressed by etomoxir, lysosomal inhibitors or rapamycin. E) Respiration of control or Plin3 silenced NIH-3T3 cells under basal condition and after 6 hours of OA treatment. Plin3 silencing prevents the increase in mitochondrial respiration. F and G) Maximal respiration and ATP production are increased in control NIH-3T3 cells after OA treatment, whilst Plin3 silencing ameliorates this effect. Bars are mean ± SEM. *p < 0.05, **p < 0.01, and ***p < 0.001 (differences caused by lysosomal inhibitors treatment), ^#^p<0.05, ^##^p<0.01, and ^###^p<0.001 (differences caused by treatment)

### mTORC1 inhibition in human liver slices drives fat accumulation

To establish the physiological relevance of our observations, excised human liver was processed to generate precision cut liver slice (PCLS) cultures in which the role of the mTORC1-Plin3 pathway could be determined in the context of intact human liver tissue ((Paish et al., 2019b), Fig. 6A). This *ex vivo* PCLS approach when coupled with use of a bespoke bioreactor technology allows the anatomical and functional maintenance of the liver tissue for over a week in culture ((Paish et al., 2019b), Fig. S4A). Liver slices were treated with OA, rapamycin and lysosomal inhibitors. Relative to untreated control PCLS, OA treatment increased autophagy activity along with enhanced LC3, p62 and Plin3 flux in the liver tissue (Fig. 6B-C, p<0.05). This increase was enhanced with rapamycin treatment (mTORC1 inhibition tested in Fig. S4B). Lysosomal degradation of Plin3 was blocked by rapamycin treatment as seen by the absence of flux (Fig. 6B-C, p<0.001). The volume of LDs was analysed by the quantification of BODIPY, this analysis showed OA treatment to increase LD volume in the liver slices (Fig. 6D, p<0.05). Consistent with the block of Plin3 degradation by rapamycin, mTORC1 inhibition also stimulated an accumulation of LDs in PCLS (Fig. 6D, p<0.001).

**Figure 6:**
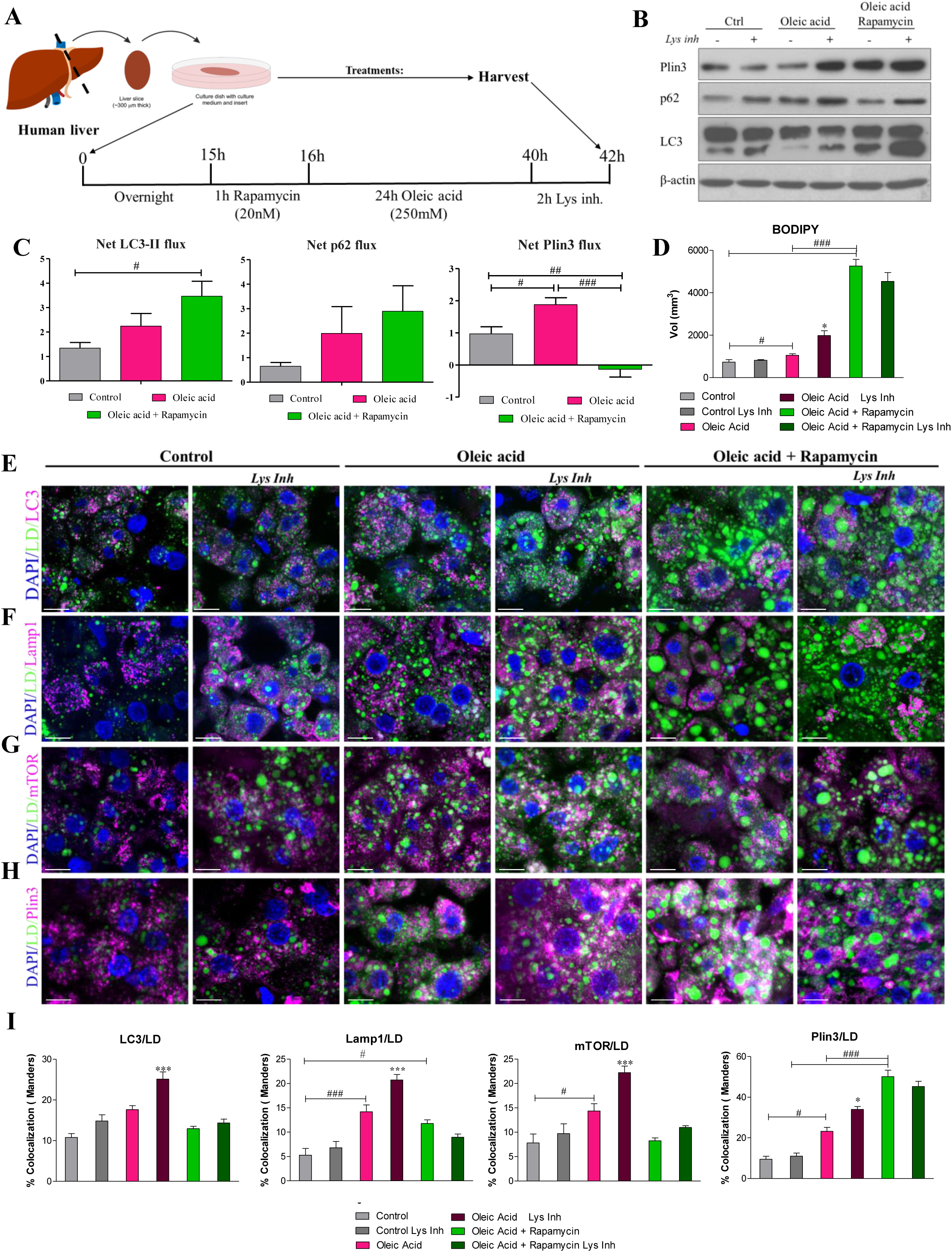
Rapamycin treatment in human liver slices drives fat accumulation. A) Depiction of the methodological approach: Human liver slices are pre-treated with or without rapamycin for 1 hour before the OA treatment, then slices are treated with or without lysosomal inhibitors. Slices without any treatment but the lysosomal inhibitors are used as a control. B-C) Levels of autophagy proteins and perilipins in total homogenates from all the conditions were analysed. Autophagy flux is elevated with the OA treatment. Rapamycin pre-treatment increases autophagy flux but leads to an accumulation of Plin3. D) Volume of LDs (measure from LD staining in green) is increased after blocking autophagy in the OA-treated slices. Rapamycin treatment results in even stronger accumulation of lipids. E-I) Recruitment of LC3 (E), Lamp1 (F) or mTOR (G) (magenta) to LDs (green) is decreased after blocking mTORC1 activity whilst Plin3 (H) (magenta) accumulates on the LD (green) surface in response to rapamycin in OA-treated human liver slices (see E to H, quantified in I). Scale bar: 20μm. Bars are mean ± SEM. *p < 0.05, **p < 0.01, and ***p < 0.001 (differences caused by lysosomal inhibitors treatment), ^#^p<0.05, ^##^p<0.01, and ^###^p<0.001 (differences caused by treatment)

Finally, we employed immunofluorescence to determine lipophagy within hepatocytes in the PCLS tissue. Colocalization of LC3 and LAMP1 with LDs was increased in hepatocytes following OA treatment. Moreover further accumulation was observed when lysosomal activity was blocked. This increased colocalization was prevented by treatment with rapamycin (Fig. 6E, F and I, p<0.05). The presence of mTOR on LDs was also increased after OA treatment, and followed the same pattern observed for the autophagic markers (Fig. 6G and I, p<0.05). Similarly, recruitment of Plin3 to LDs was enhanced following OA treatment which was further increased by lysosomal inhibitors (Fig. 6H and I, p<0.05). Importantly, rapamycin treatment promoted a strong accumulation of Plin3 on LDs (Fig. 6H and I, p<0.001). Together, these results support our observations from studies in monocultures and indicate that mTORC1-Plin3 axis plays a key role in the process of lipophagy in human liver.

## DISCUSSION

Plin3 is important in LD maturation (Bulankina, Deggerich, Wenzel, Mutenda, Wittmann, Rudolph, Burger and Honing, 2009) and may be a biomarker for the early stages of NAFLD (Pawella, Hashani, Eiteneuer, Renner, Bartenschlager, Schirmacher and Straub, 2014). Remarkably, we report here that Plin3 is a crucial regulator of lipophagy (Fig. 7). Specifically, Plin3 is required for the recruitment and assembly of protein components of the autophagy machinery to LD in response to lipid loading. Whilst Plin3 does not contain a putative LIR motif to mediate its interaction with LC3 (Jacomin, Samavedam, Promponas and Nezis, 2016), other non-canonical autophagy receptors have recently been described. For example, CCPG1 binds to FIP200 instead of LC3 to promote ER-Phagy in the pancreas (Smith and Wilkinson, 2018, Smith, Harley, Kemp, Wills, Lee, Arends, von Kriegsheim, Behrends and Wilkinson, 2018). Similarly, our Plin3 pulldowns revealed its strong association with autophagy initiator proteins, FIP200 and ATG16l, after OA treatment (Fig. 2I, J). FIP200 is a crucial player in the initiation of autophagy and, together with Atg16L1, may also be involved in later stages of autophagosome formation (Nishimura, Kaizuka, Cadwell, Sahani, Saitoh, Akira, Virgin and Mizushima, 2013). Interaction of FIP200 with Beclin1 was also affected by Plin3 silencing, further supporting its importance for autophagic degradation of LDs (Russell et al., 2013, Khaminets, Behl and Dikic, 2016, Hara, Takamura, Kishi, Iemura, Natsume, Guan and Mizushima, 2008). We also report that the ability of Plin3 to regulate the assembly of autophagic complexes on LD is dependent upon mTORC1 regulated phosphorylation. *In silico* analysis supports mTOR as a candidate kinase for Plin3 phosphorylation (http://www.phosphonet.ca/), as does our own data showing that treatment with rapamycin inhibits Plin3 phosphorylation. Of note, mTOR is hyperactivated during overfeeding ((Zoncu, Efeyan and Sabatini, 2011), Fig. S1C). Moreover, phosphorylated mTOR is found in LDs, together with the key proteins required for mTORC1 activation. Therefore, the mTORC1 pathway is active on LDs and through its regulation of Plin3 phosphorylation defines a novel cellular function for mTORC1 in lipid metabolism and energy production.

**Figure 7:**
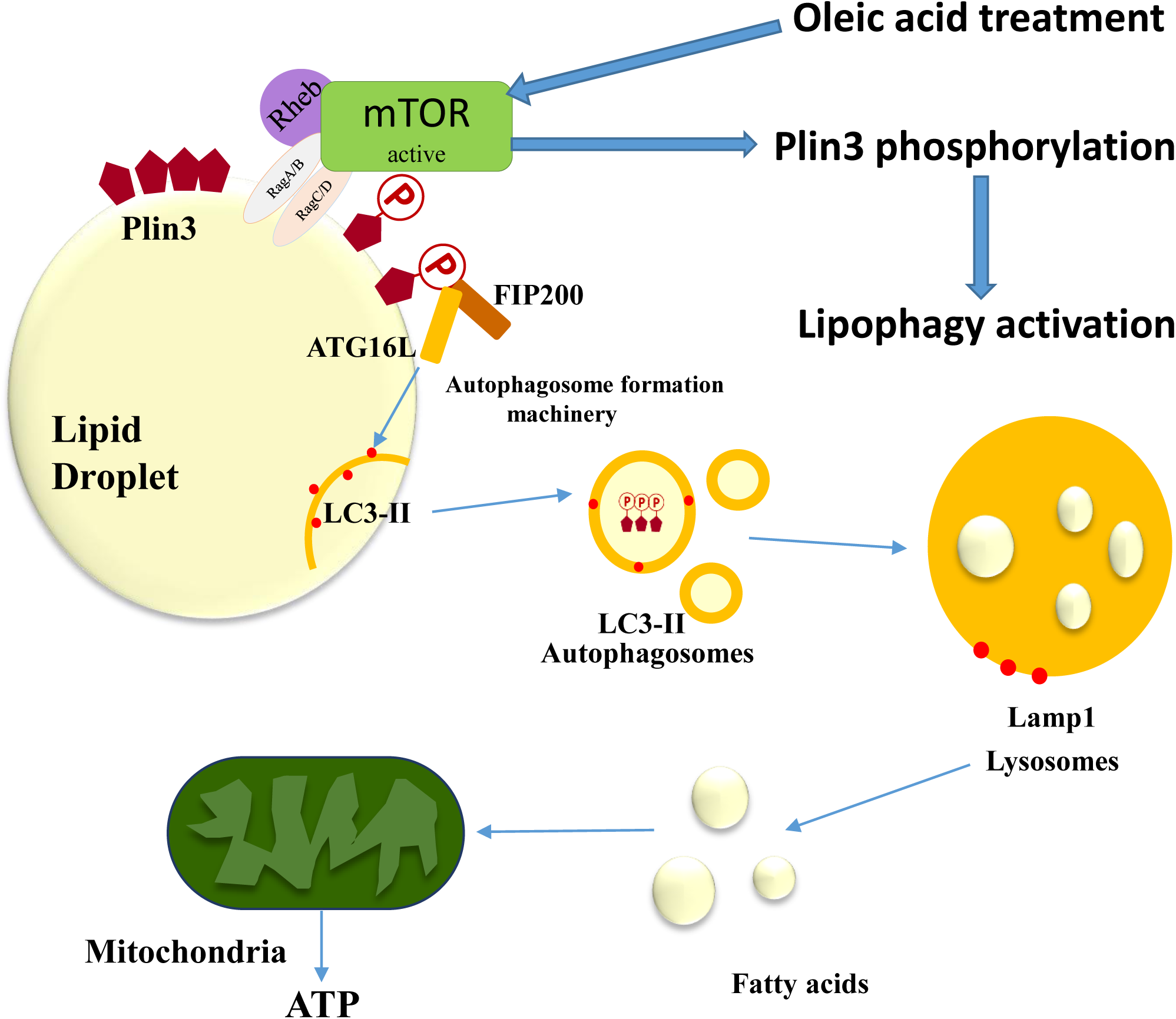
Proposed model where phosphorylated Plin3 acts as a docking protein for lipophagy. Oleic acid treatment activates mTORC1 activity to directly or indirectly phosphorylate Plin3 at LDs. Phosphorylated Plin3 binds the autophagosome formation machinery (FIP200 and ATG16l) to induce lipophagy. The autophagic machinery mobilises LDs to lysosomes, which generate fatty acids used for energy production by the mitochondria.

Clinical studies with mTORC1 inhibitors have identified several adverse side effects such as hyperlipidemia (hypertriglyceridemia and hypercholesterolemia) and activation of gluconeogenesis in the liver (Houde, Brule, Festuccia, Blanchard, Bellmann, Deshaies and Marette, 2010, Levy et al., 2006). mTORC1 is implictated in lipogenesis (Han et al., 2015) and diet-induced hepatic steatosis in mice (Kim, Qiang, Hayden, Sparling, Purcell and Pajvani, 2016), however its precise contribution to hepatic lipid metabolism remains to be fully elucidated. Although rapamycin treatment has been shown to reduce lipid hepatic synthesis (Houde, Brule, Festuccia, Blanchard, Bellmann, Deshaies and Marette, 2010), it also impairs lipid storage and catabolism in adipose tissue (Houde, Brule, Festuccia, Blanchard, Bellmann, Deshaies and Marette, 2010). Thus, lipid disbalance caused by mTORC1 inhibition is more likely to be due to defective lipid degradation rather than increased synthesis. Our experiments using rapamycin demonstrate that inhibition of mTORC1 does not result in increased autophagic degradation of LDs. Although rapamycin increased autophagic flux in the total homogenates, it prevented the interaction between Plin3 and FIP200 and ATG16l on LDs. As a result, recruitment of autophagic machinery, including autophagy initiator proteins (FIP200, Beclin1, ATG16l) as well as general autophagosome (LC3) and lysosome (Lamp1) markers, to the LD surface was suppressed by mTORC1 inhibition. Therefore, the novel mTORC1 function identified in our study may help to explain hyperlipidemia observed in response to mTORC1 inhibitors in clinical trials. Our demonstration that the mTORC1-Plin3 pathway is active in human PCLS and is required for lipophagy offers an exciting *ex-vivo* model with which to further interrogate the role of mTORC-1-Plin3 in hepatosteatosis and to determine its potential as a novel target for the prevention of NAFLD.

## Acknowledgments

We are members of the SEBBM, SEFAGIA and INEUROPA network. This work was supported by: C0120R3166, C0245R4032 and BH182173. M G-M is a Sara Borrell Postdoctoral fellow from the Ministerio de Ciencia, Innovación y Universidades (Spain). D A M, F O, L B and J M are funded by the UK Medical Research Council (Grants MR/K10019494/1, MK/K001949/1 and MR/R023026/1), National Institute on Alcohol Abuse and Alcoholism (NIAAA) (grant UO1AA018663) and the National Institute for Health Research Newcastle Biomedical Research Centre based at Newcastle Hospitals NHS Foundation Trust and Newcastle University. D A M is also funded by a CRUK Programme Grant (C18342/A23390). V I K acknowledges support from Biotechnology and Biological Sciences Research Council (BB/M023389/1, BB/R008167/1, BB/R506345/1).

## Author contributions

M G-M performed experiments and analysed the data, A S-L, J L, H P and A W assisted with the experiments. L B, J M and F O provided intellectual input. M G-M conceived the idea, designed the experiments, interpreted data and wrote the manuscript. A S-L helped to write the manuscript. V I K and D A M supervised the project and edited the manuscript. All authors commented and reviewed on the manuscript.

The authors declare no competing interests.

## MATERIAL AND METHODS

### Mice

Mice were used for hepatocyte isolation. All animal experiments were approved by the Newcastle Ethical Review Committee and performed under a UK Home Office licence (P3F79C606) in accordance with the ARRIVE guidelines. All mice were house in pathogen-free conditions and kept under standard conditions with a 12-hour day/night cycle and access to food and water ad libitum. Power calculations were not routinely performed; however, animal numbers were chosen to reflect the expected magnitude of response taking into account the variability observed in previous experiments.

### Cells culture and Hepatocyte isolation

NIH/3T3 cells were cultured in high glucose (4.5 g/l), glutamine-supplemented Dulbecco’s Modified Eagle’s Medium (DMEM) with 10% Fetal Bovine Serum and 1% penicillin/streptomycin (P/S).

Hepatocytes were isolated from C57BL/6J mice using a two-step perfusion method (Jack Leslie, 2019). Under terminal anaesthesia, mice underwent a laparotomy and cannulation of the inferior vena cava followed by clamping of the superior vena cava to achieve retro-perfusion of the liver using the portal vein as an outlet. *In situ* liver digestion was performed using collagenase from Clostridium histolyticum. Hepatocytes were isolated by tearing and agitating the perfused liver in Krebs-ringer buffer and filtering through a 70-μm cell strainer. Hepatocytes were collected by three rounds of centrifugation (50g for 3 minutes) followed by washes in Krebs-Ringer buffer. The hepatocyte rich fraction was then purified using a 40% Percoll density gradient (250g for 6 minutes). Pelleted liver hepatocytes were collected and culture in 10% FCS Williams E for subsequent experiments.

### Isolation of LD Fractions

For the isolation of LDs from cultured cells, a modified method of Brasaemle (Brasaemle and Wolins, 2006) was used. Cells were homogenized in lipid droplet (LD) buffer (20mM Tris/Cl at pH 7.4 and 1mM EDTA with phosphatase and protease inhibitors) and centrifuged at 1000g for 10min at 4C. Supernatants were mixed with OptiPrep to obtain a suspension of 30%, which was placed in ultracentrifugation tubes (SW40 tubes; Beckman Coulter GmbH, Germany). This bottom layer was overlaid with 20% and 10% Optiprep mixtures in LD buffer and finally with LD buffer supplemented with phosphatase and protease inhibitors. The gradients were centrifugated for 3 h at 4°C and 40.000rpm (SW40TI rotor). LD layers on top of the gradient were collected. This fraction was delipidated using methanol washes and solubilized in 2% SDS for immunoblotting. GAPDH was used to exclude cytosolic contamination and ponceau was used as a loading control (Martinez-Lopez, Garcia-Macia, Sahu, Athonvarangkul, Liebling, Merlo, Cecconi, Schwartz and Singh, 2016)

### Human Biopsies

Human liver tissue from surgical resections were obtained under full ethical approval (H10/H0906/41) and used subject to patients written consent. Liver cohort consisted of normal control human liver tissue, which was collected from patients undergoing surgical resections.

### Precision Cut Slices (Paish et al., 2019a)

Liver tissue was cored using an 8mm Stiefel biopsy punch (SmithKline Beecham, UK). Cores were then transferred to a metal mould and submerged in 3% low gelling temperature agarose and then cooled on ice. Agarose embedded tissue cores were super-glued to the vibratome mounting stage, submersed in the media chamber containing 4°C Phosphate buffered Saline and cut using a Leica VT1200S vibrating blade microtome (Leica Biosystems, UK) at a speed 0.3mm/sec, amplitude 2mm and thickness (step size) of 250μm. Tissue slices were transferred onto 8µm-pore Transwell-inserts and cultured in BioR plates and rocked on the bioreactor platform (patent PCT/GB2016/053310) at a flow rate of 18.136 µl/sec. Liver slices were cultured in Williams Medium E supplemented with 1% penicillin/streptomycin and L-glutamine, 1x Insulin Transferrin-Selenium X and 2% fetal bovine serum, 100nM dexamethasome 37°C supplemented with 5% CO2 and media were changed daily.

### Oleic acid treatment. Drug treatments

Cells were treated for 6 hr with 0.25 mM oleic acid (OA) to induce the formation of droplets LD (Rohwedder, Zhang, Rudge and Wakelam, 2014, Martinez-Lopez, Garcia-Macia, Sahu, Athonvarangkul, Liebling, Merlo, Cecconi, Schwartz and Singh, 2016).

To measure the effect of mTORC1 inhibition on lipophagy, cells were pretreated with Rapamycin (20μM) prior to oleic acid treatment for 1h.

To test mitochondrial use of lipids, cell were be pretreated for 12 hours with Etomoxir (CPT1a inhibitor) (100μM).

### Reagents

Antibodies for Atg16l (PM040) from MBL; Atg7 (2631), Beclin 1 (3495), FIP200 (12436), LC3B (2775), mTOR (2983), phopho-mTOR (5536), Rag A (4357), Rag C (3360), Raptor (2280), Rictor (2114), S6 (2217), phopho-S6 (4858), TSC1 (6935) and anti-rabbit (7074) were from Cell Signaling Technology; RagB (NBP1-85801) from Novus Biologicals; Plin3 (3883,for WB) from ProSci; LAMP1 (1D4B, for WB) from Developmental Studies Hybridoma Bank; PLIN3 (GP30 for CoIP) from ProGen Biotechnik; β-actin (A5441) and anti-mouse (AP130P) and anti-rat (AP136P) from SIGMA; GAPDH (ab8245), Lamp1 (ab24170 for IF) and normal IgG (ab188776) from Abcam; Fluorescence secondary antibodies (Alexa Fluor 488 and/or Alexa Fluor 647 conjugated) and anti-guinea pig (A18775) from Fisher Scientific and Beclin 1-HRP(sc-48341 HRP) and Plin 3-HRP (sc-390968 HRP) form Santa Cruz Biotechnology. The following chemicals were used: Agarose, Antimycin, Collagenase from Clostridium histolyticum, Etomoxir, FCCP, Oleic acid, Oligomycin, OptiPrep, Phosphatase and Protease Inhibitors, Protein A and Protein G agarose beads, Rotenone and Williams E Media from Sigma-Aldrich (Dorset, UK). Ammonium chloride, HCS LipidTOX, Leupeptin hemisulfate, Phospho-Tag (Wako), Pierce ECL Western Blotting Substrate and ProLong Gold antifade reagent with DAPI from Fisher Scientific (Loughborough, UK). Glutamine-supplemented Dulbecco’s Modified Eagle’s Medium (DMEM), Fetal Bovine Serum and penicillin/streptomycin (P/S) from Gibco (bought through Fisher Scientific). Bio-Rad DC Protein Assay from Bio-Rad (Hercules, CA, USA). Dexamethasome from Cerilliant. Human Albumin’s ELISA from Bethyl laboratories (Cambridge, UK)). Percoll from GE Healthcare, Life Sciences(Buckinghamshire, UK). Rapamycin from Selleck Chemicals (Houston, TX, USA).

### SDS-PAGE and immunoblotting (WB)

Whole-cell extracts were prepared in lysis buffer (20mM Tris-HCl pH 7.5, 150mM NaCl, 1mM EDTA, 1mM EGTA, 1% Triton X-100) and protein concentration of samples was determined using Bio-Rad DC Protein Assay. Proteins were separated on 8%, 10% or 12 % SDS polyacrylamide gel and transferred onto a nitrocellulose membrane in buffer containing 25 mM Tris, 192 mM glycine, and 20% methanol. Phos-tag PAGE was performed according to manufacturer’s instructions (Wako) using 100 μM Phos-tag. Blots were blocked with TBS/Tween 20 (0.1% T-TBS) containing 5% milk protein before overnight incubation with primary antibodies. Secondary HRP-conjugate antibody was use at 1:5000 dilution for an hour. The antibody complexes were detected by chemiluminescence using Pierce ECL Western Blotting Substrate. Densitometric quantification was performed on unsaturated images using ImageJ. Negative controls were performed with either no primary or no secondary antibodies. No bands were detected in any case. The results were calculated from at least three separate experiments for each antibody and were normalized to actin.

### Autophagic measurements

Flux assays were used to quantify autophagy activity. The accumulation of autophagy substrates, LC3-II or p62, or other substrates such as Plin3, in the presence of inhibitors of lysosomal proteolysis (ammonium chloride (20 mM) and leupeptin (100 μM)) reflects autophagy activity. Briefly, cells were cultured in the presence or absence of lysosomal inhibitors for 2 h (4 hours after OA treatment) following which, cells were collected and lysed and subjected to immunoblotting for LC3-II or p62. Autophagy flux was determined by subtracting the densitometric value of inhibitor-untreated LC3-II or p62 from corresponding inhibitor-treated values.

### Fluorescence microscopy

Cells were fixed on coverslips with a 4% paraformaldehyde (PFA) solution and blocked and incubated with primary and secondary antibodies (Alexa Fluor 488 and/or Alexa Fluor 647 conjugated). For lipid droplet (LD) detection, cells were incubated with BODIPY 493/503 for 20min at RT. Mounting medium (ProLong Gold antifade reagent) with DAPI (4’, 6-diamidino-2-phenylindole) to visualize the nucleus. Negative controls were performed with either no primary or no secondary antibodies. No staining was detected in any case. Images were acquired on a Leica DMi8 wide field fluorescence (Inverted) (Leica Microsystems, Germany) using X63 or X100 objective/1.4 numerical aperture. Images were acquired at the same exposure times in the same imaging session. Image slices of 0.2µm thickness were acquired and deconvolved using the Huygens (Huygens Essential, The Netherlands) acquisition/analysis software.

Liver Precision Cut Slices (PCS) were fixed overnight in 4% paraformaldehyde (PFA) solution at 4°C. After three washes in PBS, liver PCS were cryoprotected first overnight at 4°C in 15% sucrose and then 4 hours at room temperature in 30% sucrose. Liver PCS were oriented and embedded in OCT and then frozen in liquid nitrogen. 10 μm sections were obtained in a cryostat. Sections were rinse in PBS and then blocked with 3% horse serum, 1% BSA in 1x PBS containing 0.4% triton X-100 for 1 hour at room temperature, then they were incubated with primary antibodies overnight at 4C. For lipid droplet (LD) detection, sections were incubated with LipidTOX 637/655 for 1 hour at RT together with the secondary antibodies. Mounting medium (ProLong Gold antifade reagent) with DAPI (4’, 6-diamidino-2-phenylindole) to visualize the nuclei. Negative controls were performed with either no primary or no secondary antibodies. No staining was detected in any case. Images were acquired on a Nikon A1R confocal inverted (Nikon Instruments INC., NY, USA) using X100 objective/1.4 numerical aperture. Images were acquired at the same exposure times in the same imaging session.

Quantification was performed after appropriate thresholding using the ImageJ software (NIH) (Schneider, Rasband and Eliceiri, 2012) in a minimum of 30 cells from at least 3 experiments. Cellular fluorescence intensity was expressed as mean integrated density as a function of individual cell size. Percentage colocalization was calculated using the JACoP plugin in single Z-stack sections of deconvolved images.

### Gene silencing with small interfering RNA (siRNA)

Cells were transiently transfected with 100 nM of siRNA constructs or a non-related siRNA (siScramble) (Sigma-Aldrich) using Lipofectamine 2000 (Invitrogen) or INTERFERin (Polyplus) (for the Plin3 silencing) for 48 hours according to the manufacturer’s protocol: cells were exposed to the silencing mix for 6 hours in the absence of serum. Silencing medium was replaced with fresh DMEM containing 10 % FBS and experiments were performed 42 hours later. The following siRNAs were used: siScramble (sense(s)) 5’- AAUUCUCCGAACGUGUCACGU-3’, (antisense (as)) 5’-ACGUGACACGUUCGGAGAAUU- 3’; siATG7 (s) 5’-CUGUGAACUUCUCUGACGU-3’, (as) 5’-ACGUCAGAGAAGUUCACAG-3’; simTOR (s) 5’-GGAUCAACCACCAGCGCUA-3’, (as) 5’-UAGCGCUGGUGGUUGAUCC-3’; and siPlin3 (s) 5’-GAGUGCUUGUGAAAUCAGA-3’, (as) 5’- UCUGAUUUCACAAGCACUC-3’. Silencing efficiency was determined by immunoblotting for the corresponding protein product.

### Immunoprecipitation

Whole cell lysates (500 μg) or lipid droplet fractions (400 μg) lysed in Co-IP lysis buffer (50 mM Tris-HCL pH 8, 150 mM NaCl, 1 mM EDTA, 1% NP40, 0.5% NaDoc and 0.1% SDS and phosphatase and protease inhibitors) were incubated overnight in rotation at 4°C with 5μg of Plin3 or 10μg of FIP200. Non-related IgG was used as negative control. Antigen-antibody conjugates were incubated for 1 hour in rotation at 4°C with a 1:1 mix of Protein A and Protein G agarose beads. All samples were then washed in Wash Buffer and bound proteins were eluted by boiling (95°C for 5 min) in 2X SDS–PAGE sample buffer. Immunoprecipitated proteins and original lysates (input) were resolved on SDS–PAGE, and membranes were probed by immunoblotting. HRP-conjutated antibodies were used when molecular weights had similar weights as the IgGs.

### Bioenergetics

Oxygen consumption rates of NIH3T3 and hepatocytes were measured in real-time in an XF96 Extracellular Flux Analyzer (Seahorse Bioscience, Billerica, MA, USA). The instrument measures the extracellular flux changes of oxygen in the medium surrounding the cells seeded in XF96-well plates.

Assay was performed on the next day of cell plating. Regular cell medium was then removed and cells were washed twice with DMEM running medium (XF assay modified supplemented with 11 mM glucose, L-glutamine 2 mM, sodium pyruvate 1 mM, pH 7.4) and incubated at 37°C without CO2 for 1 h to allow cells to pre-equilibrate with the assay medium. Oligomycin, FCCP or antimycin/rotenone diluted in DMEM running medium were loaded into port-B, port-C or port-D, respectively. Final concentrations in XF96 cell culture microplates were 1.5 μM oligomycin, 20 μM FCCP and 2.5 μM antimycin and 1.25 μM rotenone. The sequence of measurements was as follow unless otherwise described. Basal level of oxygen consumption rate (OCR) was measured 5 times, and then port-A was injected and mixed for 3 minutes, after OCR was measured 3 times for 3 minutes. Same protocol with port-B and port-C. OCR was measured after each injection to determine mitochondrial or non-mitochondrial contribution to OCR. All measurements were normalized to average three measurements of the basal (starting) level of cellular OCR of each well. Each sample was measured in 3–5 wells. Experiments were repeated 3–5 times with different cell preps. Non-mitochondrial OCR was determined by OCR after antimycin/rotenone injection. Maximal respiration was determined by maximum OCR rate after FCCP injection minus non-mitochondrial OCR. ATP production was determined by the last OCR measurement before oligomycin injection minus the minimum OCR measurement after oligomycin injection.

### Enzyme-linked immunosorbent assay (ELISA)

Media samples collected from precision cut human liver slices before and after each treatment. Then, ELISA quantifications for human albumin (E88-129) were performed as per manufacturer’s instructions (Paish et al., 2019a).

### Statistical Analysis

Results are presented as means ± SEM. Graphpad prism was used to perform analysis of variance (ANOVA) with a Tukey’s post hoc test. * P<0.05, ** P<0.01 or *** P<0.001 and ^#^p<0.05, ^##^p<0.01, and ^###^p<0.001 was considered statistically significant.

## SUPPLEMENTARY FIGURES

**Supplementary figure S1**: A) Levels of autophagy proteins and perilipins in primary hepatocytes after 4 or 6 hour treated with OA. B) Atg7 expression in control and Atg7 silenced NIH-3T3 cells. C) Levels of Plin3 in control and Plin3-silenced NIH-3T3 cells.

**Supplementary figure S2**: A) Levels of phospho-S6 and total S6 in primary hepatocytes before and after OA treatment. B) Levels of phospho-S6 and total S6 in primary hepatocytes without and with rapamycin treatment and without and with OA treatment. C) Levels of mTOR, phospho-S6 and total S6 in control and mTOR-silenced NIH-3T3 cells with and without OA treatment. D-G) Recruitment of LC3 and Lamp1 (magenta) to LDs (green) is decreased after silencing mTOR (quantified in E and G). H-I) mTOR silencing causes Plin3 (magenta) accumulation on LDs (green) (quantified in I).

Scale bar: 20μm. Bars are mean ± SEM. *p < 0.05 (differences caused by lysosomal inhibitors treatment).

**Supplementary figure S3:** A) Mitochondrial respiration in control or Plin3-silenced NIH-3T3 cells under basal condition or 6 hours after OA treatment with and without etomoxir (100nM), rapamycin or lysosomal inhibitors pretreatment. B and C) Maximal respiration and ATP production in control or Plin3 silenced NIH-3T3 cells in all the above conditions.

Bars are mean ± SEM. *p < 0.05, **p < 0.01, and ***p < 0.001 (differences caused by lysosomal inhibitors treatment), ^#^p<0.05, ^##^p<0.01, and ^###^p<0.001 (differences caused by treatment).

**Supplementary figure S4**: A) Soluble albumin levels in the media of human liver slices from the experimental timepoints and treatments (OA, rapamycin and lysosomal inhibitors) used in this study. B) Levels of phospho-S6 and total S6 in liver slices without and with rapamycin treatment and without and with OA treatment.

