## Supplemental figures for "mTORC1-Plin3 pathway is essential to activate lipophagy and protects against hepatosteatosis"

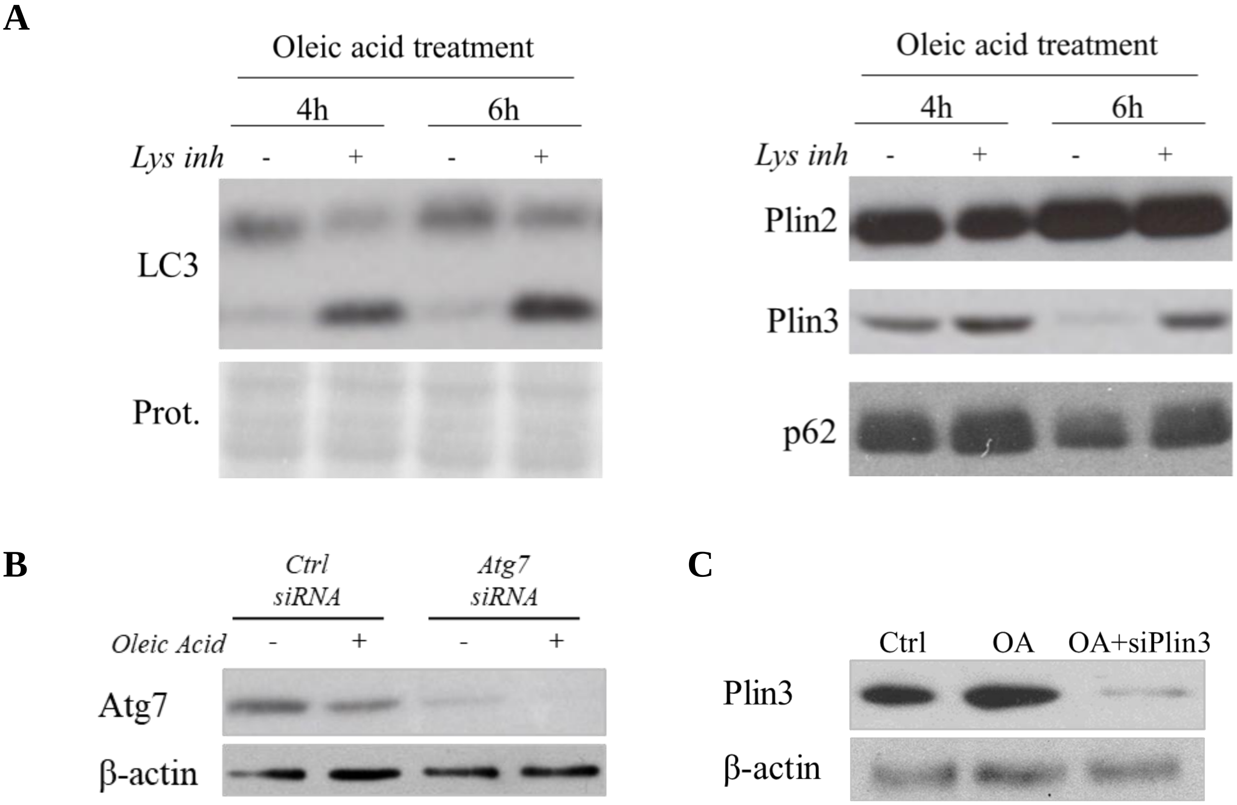


Garcia-Macia M et al. Figure S1

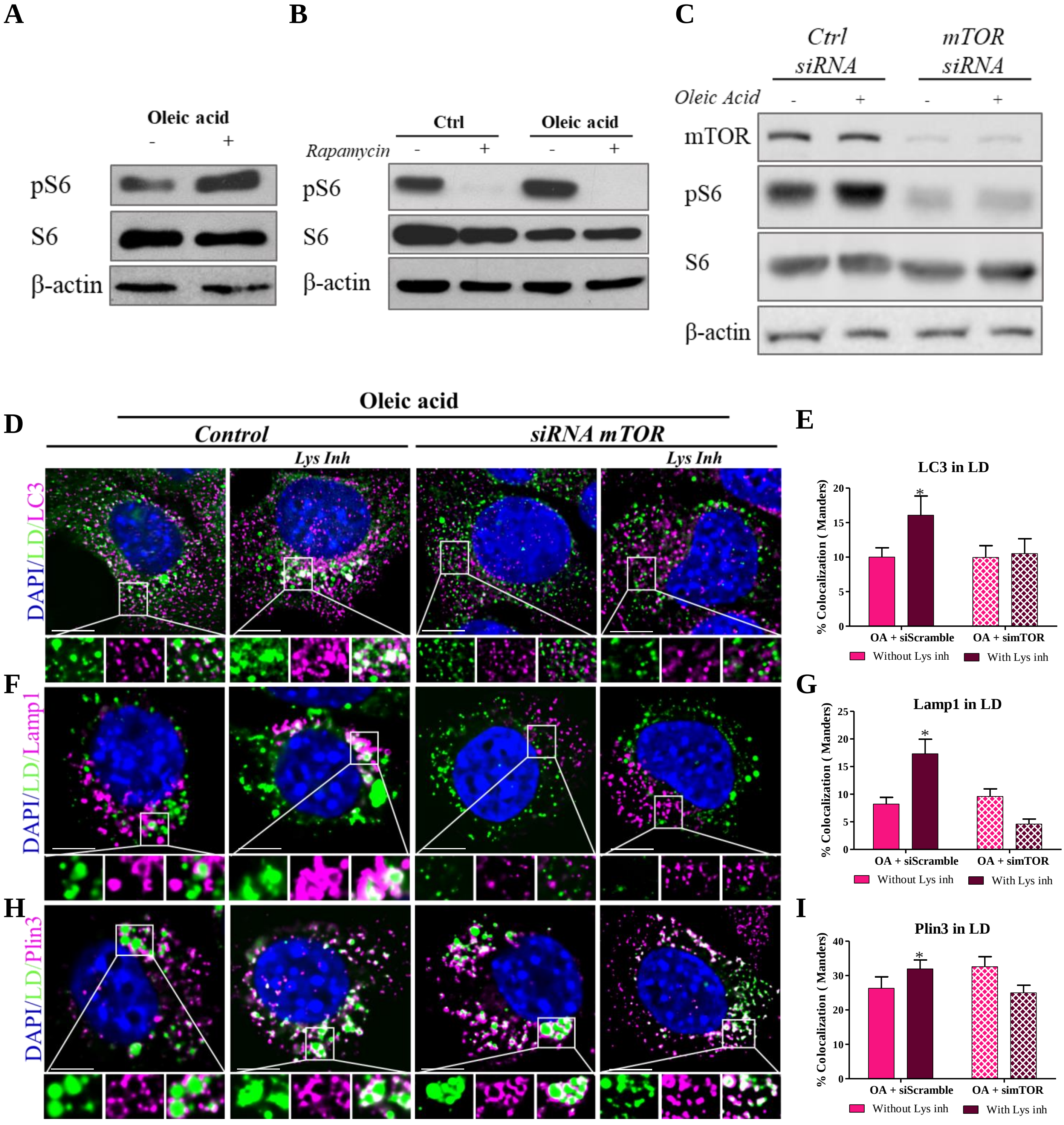


Garcia-Macia M et al. Figure S2

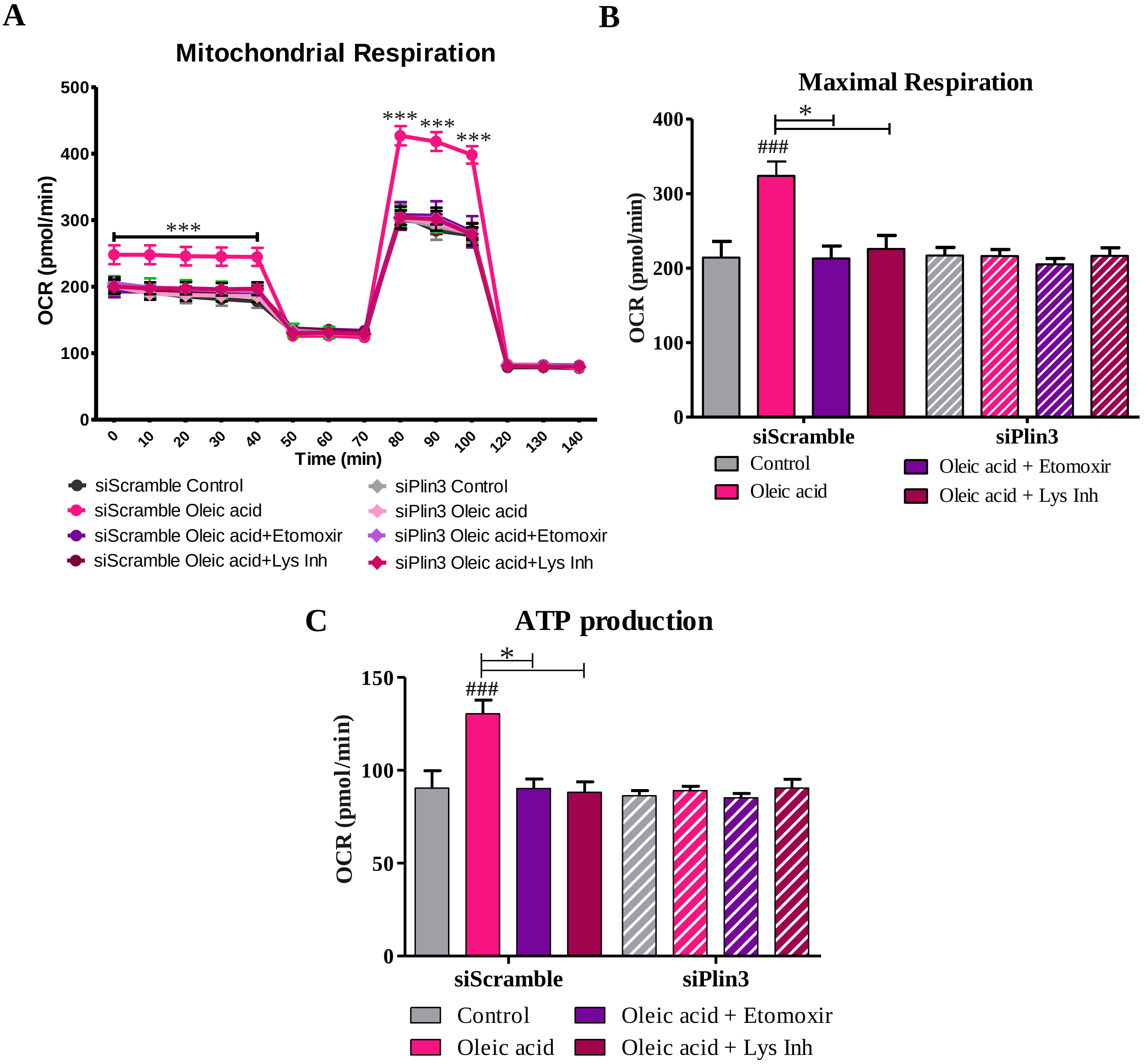


Garcia-Macia M et al. Figure S3

A

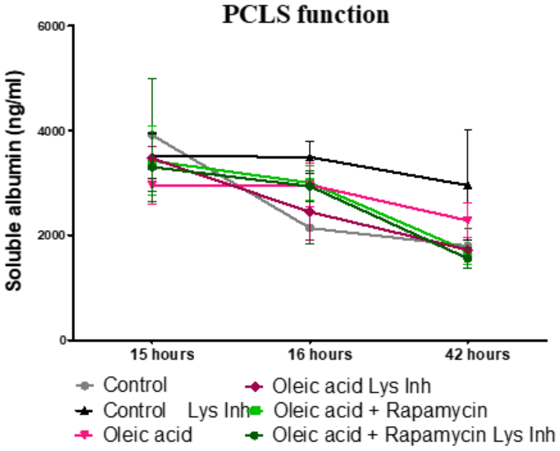


B

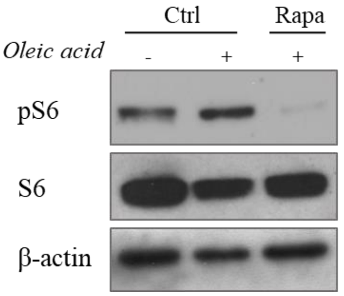


**Garcia-Macia M et al. Figure S4**
